# Caecal dysfunction in the NL3^R451C^ mouse model of autism

**DOI:** 10.1101/2022.06.15.494637

**Authors:** Chalystha Yie Qin Lee, Gayathri K. Balasuriya, Madushani Herath, Ashley E. Franks, Elisa L. Hill-Yardin

## Abstract

The mouse caecum is a pouch-like structure that is anatomically similar to the human appendix and is hypothesised to serve as a reservoir for commensal bacteria. The gastrointestinal tract is also home to the largest immunological organ of the body and the enteric nervous system (ENS), which regulates gut motility and secretion. The caecum is therefore an ideal location to study neuro-immune-microbe interactions in gut-brain communication. Individuals with Autism Spectrum Disorder (ASD; autism) frequently present with gastrointestinal symptoms in addition to core diagnostic behavioural features, implying a gut-brain link. More broadly, changes in gut-brain connectivity are now thought to play a critical role in a range of neurodevelopmental disorders. Here, we employed a mouse model of autism expressing a missense mutation in the neuroligin-3 post-synaptic protein that affects brain and enteric neuronal activity (NL3^R451C^ mice). We previously observed abnormal caecal ENS architecture and immune cell morphology in the caecal patch in this model, however it is unknown if caecal function is altered in NL3^R451C^ mice. Using a tri-cannulation approach to record motility patterns in the mouse caecum, we identified novel caecal motor complexes in *ex vivo* preparations. Caecal permeability and neurally-evoked secretion levels were also studied. Key immune populations including gut macrophages and dendritic cells within the caecal patch were stained using immunofluorescence to investigate shifts in immune activity. Caecal motility patterns in NL3^R451C^ mice differed from wildtype littermates. Specifically, caecal motor complexes occurred at a higher frequency and for a shorter duration in NL3^R451C^ mice than in wildtype littermates. In NL3^R451C^ mice, neurally-evoked caecal secretion was reduced in response to the nicotinic acetylcholine receptor agonist (DMPP), but permeability was unchanged. Increased numbers of caecal patches were observed in NL3^R451C^ mice compared to wildtype, with no alterations in morphology of selected immune populations. Future research is warranted to better understand caecal function and how neuro-immune interactions in the caecum affect health and influence GI function in neurodevelopmental disorders via the gut-brain axis.

## Introduction

The mouse caecum is synonymous with the human appendix which is widely and incorrectly known to be a vestigial remnant in the human digestive tract (Girard-Madoux et al., 2018; Smith et al., 2009). The human appendix is hypothesised to function as a reservoir for commensal bacteria (Bollinger et al., 2007; Laurin et al., 2011). The caecum is located at the junction of the distal ileum and proximal colon. The caecal environment is enriched with biofilms and mucus that support the growth and maintenance of gut flora. This allows for the re-inoculation of the colon should contents be purged via diarrhoea resulting from contact with a pathogen (Bollinger et al., 2007). The caecum also plays an immunological role via the activity of the gut-associated lymphoid tissue (GALT) located at the blind end of the pouch-like structure, termed the caecal patch. It is worth noting that in humans, the appendix is also rich in lymphoid tissue that is involved in antibody development and the maturation of T and B lymphocytes (Girard-Madoux et al., 2018; Kooij et al., 2016; Vitetta et al., 2019). The caecal patch regulates IgA secretion that modulates immune selectivity; in basic terms, this role assists in determining if bacteria are friend (commensal) or foe (pathogen) (Kooij et al., 2016; Masahata et al., 2014). Within the gut wall in close proximity to the GALT and the microbial reservoir, the enteric nervous system (ENS) is present, providing an ideal location for immune-neuronal crosstalk (Lee et al., 2021). The ENS is composed of two neuronal ganglia (the submucosal and myenteric plexus) that primarily function to regulate gut motility and permeability, respectively. The caecum is a targeted region of interest due to the presence of multiple key components regulating neuro-immune-microbe interactions in close proximity to one another. Although its specific role remains to be clarified, the caecum is therefore also ideally located as a potential ‘organising node’ of the gastrointestinal tract (Lee et al., 2021). Despite mounting interest in the role of the gut-brain axis, details of caecal motility and intestinal barrier function, which feasibly influence caecal immune and microbial function in mice, are lacking. Furthermore, techniques to examine for differences in caecal function such as motility have not been available to date.

Alterations in gut-brain communication pathways are now widely implicated in many neurological disorders. Neurodevelopmental disorders are characterised by cognitive, behavioural, and/or physical impairments caused by aberrant brain development. Autism Spectrum Disorder (ASD; autism) is one of the most common neurodevelopmental disorders and affects approximately one in every 44 children (Maenner et al., 2021). ASD is clinically defined by stereotyped behaviours as well as difficulties in social interaction and communication (American Psychiatric Association & DSM-5 Task Force, 2013). In addition to these characteristics, children with ASD are more likely to suffer from gastrointestinal problems. Children with autism experience general gastrointestinal (GI) illness 4 to 8 times more frequently compared to neurotypical children (Chaidez et al., 2014; Kohane et al., 2012; McElhanon et al., 2014; Schieve et al., 2012). This group has a higher occurrence of constipation, diarrhoea, and stomach discomfort than the general population. These GI symptoms are closely associated with the severity of autism, as well as increased irritability, anxiety, and social disengagement (Gorrindo et al., 2012).

Individuals with ASD also tend to have increased intestinal permeability, commonly referred to as “leaky gut syndrome”. This is based on findings from lactulose/mannitol testing (de Magistris et al., 2010), serum zonulin levels (where zonulin is a barrier protein that influences GI permeability) (Esnafoglu et al., 2017), and *post-mortem* analyses of critical intestinal barrier proteins (Fiorentino et al., 2016). These alterations could be caused by modification of tight junction proteins in the GI tract which are crucial for supporting mucosal barrier integrity (de Magistris et al., 2010; Esnafoglu et al., 2017; Fiorentino et al., 2016). When this barrier is compromised, bacteria and metabolites can potentially penetrate the GI wall and trigger excessive immune responses. High levels of inflammatory cytokines have been reported in children with ASD and this finding was additionally correlated to symptom severity (Ashwood et al., 2011; Masi et al., 2017). Altered levels of inflammatory cytokines are also linked with key ASD-associated bacterial populations (e.g., Clostridiales) and GI symptoms (Luna et al., 2017). Furthermore, chronic neuroinflammation, altered neuronal structures and cytokine profiles as well as GI immune dysfunction are prevalent in both individuals diagnosed with ASD and preclinical models of autism (Matta et al., 2019). These findings indicate that neuro-immune changes are associated with autism.

Although autism is traditionally considered to be a central nervous system (CNS) disorder, the observation of gastrointestinal symptoms highlights the potential influence of the gut-brain axis. To study the complex connection between the gut and brain in a controlled manner, the use of animal models in research is vital. The NL3^R451C^ mouse model is a well-validated animal model for studying autism. These mice are genetically engineered to possess the human Arg to Cys substitution at position 451 of the Neuroligin 3 protein (NL3^R451C^). This mutation was first discovered in two Swedish brothers diagnosed with ASD (Jamain et al., 2003), who also displayed prominent gastrointestinal symptoms (Hosie et al., 2019). Although ASD is inherently a human behavioural disorder and difficult to model in animals, mice with this mutation demonstrated attentional abnormalities, heightened repetitive behaviours, altered social and behavioural interactions, impaired spatial learning, atypical aggressive behaviours, and abnormal mating behaviours compared to wildtype littermates (Burrows et al., 2017; Burrows et al., 2015; Burrows et al., 2021; Etherton et al., 2011; Hosie et al., 2018; Jaramillo et al., 2014; Lee et al., 2020; Rothwell et al., 2014; Tabuchi et al., 2007). NL3^R451C^ mice also display imbalanced excitatory and inhibitory neuronal activity indicative of altered synaptic neurotransmission which may contribute to the abnormal behaviours observed in this model (Etherton et al., 2011; Földy et al., 2013; Hill-Yardin et al., 2015; Hosie et al., 2018; Medina et al., 2018; Tabuchi et al., 2007).

Clinical records of the two brothers expressing the R451C mutation reported GI symptoms including oesophageal regurgitation, chronic gut pain and diarrhoea (Hosie et al., 2019). Thus, gut dysfunction in the NL3^R451C^ mouse model of autism was also investigated. NL3^R451C^ mice displayed faster *in vivo* small intestinal transit and *ex vivo* colonic motility assays show increased sensitivity to GABAA receptor modulation (Hosie et al., 2019). The proportion of enteric neuronal subpopulations in the ENS are also altered in these mice. Specifically, the number of myenteric neurons per ganglia and nitric oxide synthase (NOS)-producing neurons are increased in the proximal jejunum of NL3^R451C^ mice (Hosie et al., 2019). NOS neurons are a key subpopulation of neurons in the ENS because they produce nitric oxide (NO), the major inhibitory neurotransmitter in the gut. Alterations in this NOS subpopulation of enteric neurons might contribute to modified gut function relevant to ASD.

Considering the heterogeneity in cellular architecture, digestive function (Bowcutt et al., 2014; Thompson et al., 2018), and immune system (Mowat & Agace, 2014) along the gastrointestinal tract, it is important to assess for region-specific differences in the gut relevant to ASD. We previously reported changes in neuronal proportions and immune cell morphology suggesting aberrant neuro-immune interactions in the caecum in NL3^R451C^ mice (Sharna et al., 2020). Interestingly, caecal weight was significantly decreased in mutant mice compared to wildtype littermates. The proportion of NOS neurons per ganglia as well as the total number of neurons per ganglia are increased in both the myenteric and submucosal plexus of the ENS in NL3^R451C^ mice (Sharna et al., 2020). Within the caecal patch, Sharna and colleagues found an increase in the density of gastrointestinal macrophages and the macrophage morphology was skewed towards an activated phenotype (Sharna et al., 2020). Here, we aimed to understand whether the digestive, immune, and barrier function of the caecum is altered in accordance with the previously observed shift in enteric neuronal subpopulations. Specifically, we studied the digestive churning function of the caecum by visualising caecal motility patterns using video imaging. We additionally quantified mucus area as well as caecal content weight to identify factors potentially contributing to the observed reduction in caecal weight. Caecal permeability and secretion were studied using the Ussing chamber system to elucidate functional effects of increased numbers of caecal submucosal neurons in NL3^R451C^ mice.

## Methods

### Animals

Adult male NL3^R451C^ mice and wildtype littermates (8-14 weeks old) were used in this study. These mice were bred and maintained on a C57BL/6J background at The University of Melbourne Animal Facility and transported to RMIT University Animal Facility at 8 weeks of age. All mice were culled by cervical dislocation in accordance with strict guidelines as approved by the RMIT Animal Ethics Committee (AEC#: 1727). Gastrointestinal tissue was immediately dissected upon culling for fresh tissue experiments or placed in fixative for downstream analysis.

### Ex vivo caecal motility assay

A novel *ex vivo* caecal motility assay was designed to visualise motility patterns of the mouse caecum in NL3^R451C^ mice and wildtype littermates. This method was adapted from a well-established and widely-used colon motility assay (Swaminathan et al., 2016) and a tri-cannulation technique that was previously reported in rabbit caecum (Hulls et al., 2012; Hulls et al., 2016). Briefly, the freshly dissected caecum was placed in an organ bath filled with warmed carbogenated Krebs physiological saline (composition: 118 mM NaCl, 4.6 mM KCl, 2.5 mM CaCl_2_, 1.2 mM MgSO_4_, 1 mM NaH_2_PO_4_, 25mM NaHCO_3_, 11 mM D-glucose; bubbled with 95% O_2_, 5% CO_2_). The tri-cannulation method was utilised to establish a stable intraluminal pressure within the caecal preparation mimicking physiological conditions. The ileum end of the tissue was connected to a reservoir filled with Krebs, and the caecal tip and colon end was connected to an outflow tube, where a front and back pressure of about 4 cm H_2_O was established. Krebs physiological saline was continuously superfused through the organ bath at a constant flow rate of 8 ml min^−1^ and the organ bath temperature was maintained at a constant 36 °C. Video recordings of caecal motility were recorded *ex vivo* on the Virtual Dub video capture software (version 1.10.4, open source) using a Logitech webcam (QuickCam Pro 4000; I-Tech, Ultimo, NSW, Australia) mounted at a fixed height directly above the organ bath. A purpose-built, in-house open-source software (Analyze2) was used to convert video files of 15-minute duration each to spatiotemporal heatmaps. The colour of each pixel on the heatmap represents the diameter of the caecal preparation, where the cooler colours (blue, green) denotes a wider diameter or relaxed gut and the warmer colours (red, yellow) denotes a narrow diameter or constricted gut. The x-axis of the heatmap represents time lapsed and the position along the length of caecal segment is represented on the y-axis. The overall experimental protocol consists of a 30-minute equilibration period followed by four 15-minute (total 1 hour) duration recordings of baseline caecal motility.

From the spatiotemporal heatmaps of caecal motility produced, several indices of neurally-mediated caecal motor activity were compared between wildtype and NL3^R451C^ mice. Specifically, we analysed caecal motor complexes (CaeMC) and gut resting diameter. CaeMCs are clusters of individual contractions that typically begin with a forward contraction (i.e., a contraction travelling from caecal tip to colon), followed by multiple reverse (that is, contractions travelling from colon to caecal tip) and forward contractions. A study of rabbit caecum motility (Hulls et al., 2012) refer to these contractions as pro and anti-peristaltic contractions, respectively. In mice, these clusters or complexes are interspersed by periods of quiescence where little to no contraction activity occurs. We note that similar nomenclature is used for colonic motor complexes (CMC), which are regular neurally-regulated contractions that propagate along the colon (Spencer et al., 2020). CaeMC frequency was measured by manually counting the number of CaeMCs observed in one heatmap (15 min). CaeMC velocity, CaeMC duration and the period of quiescence between each CaeMC were measured by annotating the region of interest using the “Annotate Contractions” feature within the Analyze2 software user interface. The velocity and duration of individual contractions within a CaeMC were also analysed using the same method. We additionally identified the proportion of CaeMCs that display the canonical forward-reverse contraction pattern observed in most CaeMCs and the time elapsed between the start of a CaeMC and the start of a contraction in the attached colon tissue segment. Resting gut diameter was measured at timepoints between CaeMC in the presence of constant intraluminal pressure. This is done by taking cross sections of the spatiotemporal heatmap at three regions corresponding to the respective caecal regions: ampulla (at the first 20% of caecal length), mid-caecum (60%), and caecal tip (80%).

### Caecal content, histology, and mucus content analysis

To obtain caecal content weight, caecal samples were first weighed before clearing. The caecum was then cut open along the mesenteric border, pinned out, and the contents flushed out using a Pasteur pipette filled with 0.1M PBS. The remaining caecal tissue was then weighed a second time and the weight difference between pre- and post-clearing was calculated as caecal content weight and normalized against mouse body weight.

For Alcian Blue staining, caecum samples were collected and immediate placed in freshly prepared methanol-Carnoy’s fixative (composition, V/V: 60% absolute methanol, 30% chloroform, 10% glacial acetic acid) overnight at 4°C. Methanol-Carnoy’s fixative preserves mucus architecture in comparison to the conventional fixation methods with 4% formaldehyde or paraformaldehyde. This was followed by two washes in absolute methanol for 30 minutes each, and two washes in absolute ethanol for 20 minutes each. After the washing steps, the tissue was processed in two rounds of xylene, 15 minutes each. The tissue was then embedded in paraffin and sectioned at 4 μm thickness. Next, the sections were stained with 1% Alcian blue in 3% Acetic Acid pH 2.5 (Sigma-Aldrich, Germany) for 10 minutes, which stains acidic mucosubstances and mucins a deep blue colour, and counterstained with Nuclear Fast Red (Sigma-Aldrich, Germany) for 5 minutes. The Olympus Slide Scanner Microscope (Olympus Australia Pty. Ltd.; Melbourne, Australia) was used to image the slides. Tissue processing and Alcian Blue staining steps were performed by the Biomedical Sciences Histology Facility of the University of Melbourne (Parkville, Australia). Mucus content in the cross section of the caecum was then quantified using FIJI ImageJ (ImageJ 1.52a, NIH) (Schindelin et al., 2012). The area of mucus stained by Alcian Blue was selected using the colour threshold feature in FIJI Image J. The number obtained was then normalized against the total area of the caecum cross section.

### Ussing Chamber permeability and secretion assays

Prior to preparing the caecal tissue for the Ussing chamber assay, the 4-chamber P2400 EasyMount Ussing Chamber system was set up using Ag/AgCl pellet electrodes for voltage sensing and Ag wire electrodes for current passing, immersed in 3 M KCl solution. The electrodes were connected to each half of the chamber (1 voltage and 1 current electrode in each compartment) by salt agar bridges (electrode tips filled with 2.5 % agarose melted in 3 M KCl solution). The electrodes were then connected to the EC-825A Epithelial Voltage Clamp (Warner Instruments, Holliston, MA, USA). The Ussing Chamber was connected to a bench-top water bath that water-jacket each chamber to ensure the temperature of Krebs solution within the chamber is maintained at 37 °C. Fluid resistance was compensated with Krebs solution in the chambers and an empty P2404 Ussing chamber slider. Asymmetries in the voltage measuring electrodes were eliminated by applying an offset voltage to nullify the voltage difference between the two voltage sensing electrodes in each compartment of the chamber. The Ussing Chamber system, voltage electrodes, current electrodes, electrode tips, and sliders were obtained from Physiologic Instruments, San Diego, CA, USA.

The caecum was dissected and immediately placed in ice-cold Krebs solution bubbling with carbogen. The sac-like structure was cut open along the inner mesenteric border using small spring scissors and pinned flat onto a black Sylgard-lined (Dow Corning, Midland, MI, USA) petri dish. The luminal contents within the caecum were gently flushed away with Krebs solution. To obtain the mucosa-submucosa layers, the pinned-out caecum was re-pinned with the mucosa side facing down. The serosa, longitudinal muscle and circular muscle layers were carefully peeled away under a stereomicroscope (Olympus, SZ61, Japan). The peeled caecum preparation was then mounted on the pins of one side of an Ussing chamber slider. The slider with the mounted tissue was then immediately placed in the slot at the middle of the chamber. The mucosal and serosal compartments are independently superfused with 5 mL of Krebs solution and constantly carbogenated throughout the experiment.

Caecum preparations were allowed to equilibrate for 20 min prior to collecting measurements. The transepithelial resistance (TER) was measured by applying a voltage-clamp of 1 mV and recording the resulting short-circuit current (I_SC_) reading. This procedure was repeated 5 times at 50 s intervals to obtain an average reading. The TER was then calculated using Ohm’s Law, whereby

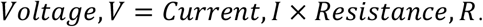

After measuring the TER, paracellular permeability of the gut tissue was assessed. Fluorescein isothiocyanate-dextran (4 kDa FITC-dextran, Sigma Aldrich, Australia) was added to the mucosal compartment by replacing 250 μL of Krebs solution with 10 mg/mL FITC-Dextran stock solution to a final concentration to 0.5 mg/mL. At each timepoints, 200 μL of solution was collected from the serosal compartment and replaced with fresh solution to ensure that equal volumes are maintained in both compartments. Samples were collected at t = 0, 15, 30, 35, 40, 45, 50, 55, 60, 70, 80, 90, 105, 120, 135 min. TER readings were also taken at 15-min intervals to assess the change in TER upon addition of FITC-Dextran into the system. Absorbance reading of the samples at each timepoint were taken on the FlexStation 3 Microplate Reader (Molecular Devices, Sunnyvale, CA, USA) (Excitation 485 nm; Emission 538 nm). FITC-Dextran concentration in the serosal compartment was then calculated based on a standard curve of known concentrations.

Electrogenic secretion of the caecal preparation was assessed by measuring short-circuit current (I_SC_) in the presence of a ganglion-specific nicotinic acetylcholine receptor (nAChR) agonist, 1,1-dimethyl-4-phenylpiperazinium (DMPP) (Prado & Segalla, 2004). The mucosal-submucosal caecal preparation was mounted in an Ussing chamber slider as previously described for TER and permeability measurements. The transepithelial voltage across the preparation was then clamped at 0 mV in voltage-clamp mode. The resulting short-circuit current (I_SC_) was monitored at baseline and when DMPP is added to the serosal compartment of the chamber.

An non-specific cholinergic agonist, carbachol was added to the serosal compartment of the chamber at the conclusion of all experiments to ensure that tissue is still viable. Tissues were discarded if there was no response (i.e., the well-established response to Carbachol application being a significant increase in I_SC_). The maximum duration of all experimental protocols was restricted to 3 hours after mounting the tissue onto the slider to ensure optimum tissue integrity throughout experiments.

### Microdissection and immunofluorescence staining

The freshly dissected caecum was cut open along the mesenteric border, stretched flat and pinned out on a Sylgard-coated (Dow Corning, Midland, MI, USA) petri dish. The preparation was fixed in 4% formaldehyde (Sigma-Aldrich, Germany) overnight at 4°C. After three 0.1 M PBS washes, the caecal patch was excised using dissection spring scissors and stored in 30% sucrose overnight at 4°C as a cryoprotective step. The caecal patch was then embedded in Tissue-Tek O.C.T. compound (Sakura Torrance, CA, USA) for cryotomy at 10 μm per section for subsequent staining. The myenteric plexus wholemount preparation was obtained by peeling the mucosal and circular muscle layers of the pinned-out caecum. Caecum preparations for obtaining submucosal plexus wholemounts were pinned and fixed in 4% paraformaldehyde (Sigma-Aldrich, Germany) prior to microdissection. To obtain the submucosal plexus layer, the mucosal layers were carefully lifted and separated from the bottom muscular layers using fine spring scissors, and the upper layer of villi were gently scraped off using the blunt edge of curved forceps. The sections and wholemount preparations were blocked and stained according to the antibodies and concentrations listed in Table 1 Below.

**Table 1:** Concentrations of blocking solutions, primary and secondary antibody steps.

| Tissue | Blocking step | Primary antibody | Secondary antibody |
| --- | --- | --- | --- |
| **Caecal patch cross-section** | 2% donkey serum /<br>0.1% Triton-X | Goat anti-IgD (1:500)<br>Rabbit anti-CD11c 647 (1:500) | Donkey anti-sheep 488 (1:400) |
| **Myenteric plexus** | 10% CASBlock /<br>0.1% Triton-X | Human anti-Hu (1:5000)<br>Sheep anti-NOS (1:1000)<br>Rabbit anti-Iba1 (1:400) | Donkey anti-human 594 (1:750)<br>Donkey anti-sheep 488 (1:400)<br>Donkey anti-rabbit 647 (1:400) |
| **Submucosal plexus** | 1% Triton-X | Human anti-Hu (1:5000)<br>Rabbit anti-VIP (1:100)<br>Goat anti-ChAT (1:100) | Donkey anti-human 594 (1:750)<br>Donkey anti-sheep 647 (1:400)<br>Donkey anti-rabbit 488 (1:400) |

### Analysis of neurons and immune cell populations

Caecal neurons from both the myenteric and submucosal plexus were analysed using Image J software (ImageJ 1.52a, NIH, Bethesda, MD, USA), where 10 ganglia from each preparation were selected and counted for their respective stained neuronal targets. Imaris software (Imaris 64X 9.1.0; Bitplane AG, UK) was used for 3D reconstruction and analysis of Iba-1^+^ and CD11c^+^ cells.

### Statistical analysis

GraphPad Prism version 9.1.2 for Windows (GraphPad Software, San Diego, CA, USA) was used to perform statistical analyses on the data obtained. CaeMC frequency from the motility assays were analysed using a Mann-Whitney statistical test. All measures of caecal motility were assessed using the Benjamini-Hochberg procedure to account for false discovery rate. The rest of the data were analysed using two-tailed unpaired *t*-tests, one-way and two-way ANOVA statistical tests accordingly. Data are presented as mean ± standard error of mean (SEM).

## Results

### Distinct motility patterns observed in the caecum

Our pseudo-coloured spatiotemporal heatmaps produced from video recordings of caecal motility show for the first time that this region of the mouse GI tract produces a novel rhythmic contractile pattern that occurs over time. These contractions observed are robust, regular, and clearly visualised in real time (**Supp. Video 1**). We show that caecal contraction patterns comprise multiple clusters of individual contractions extending from the caecal tip. These motility patterns appear to correlate with contractile activity in the proximal region of the colon (which is attached in the experimental preparation we examined). Each cluster includes multiple forward (from caecal tip to colon) and reverse (from colon to caecal tip) contractions (**Fig 1**). Here, we refer to these clusters of contractions as Caecal Motor Complexes (CaeMCs) (**Fig 1B** wildtype, **1D** NL3). In mice, each contractile complex is comprised of multiple consecutive forward and reverse contractions in varying order (**Fig 1B** wildtype, **1D** NL3).

**Figure 1:**
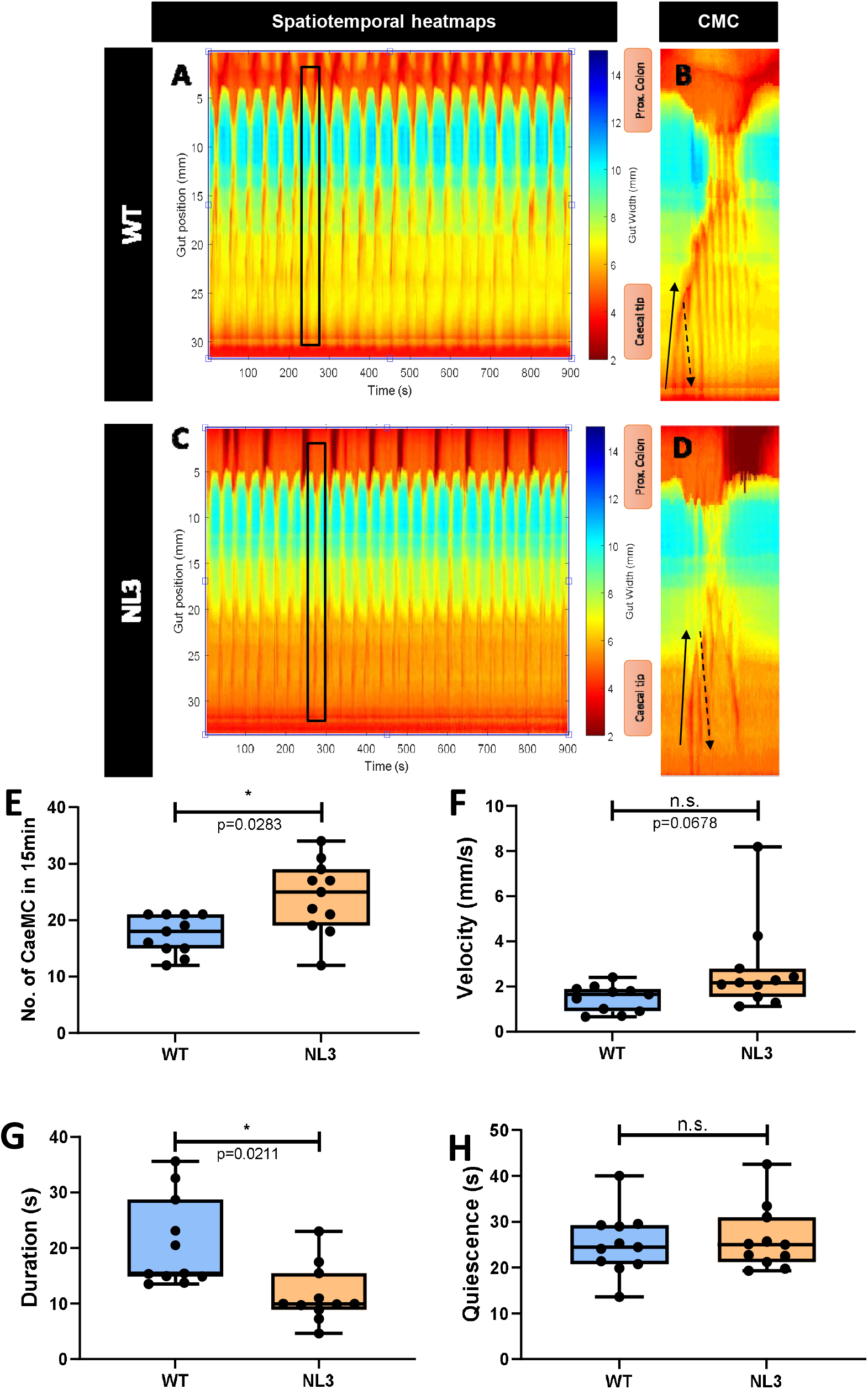
Caecal motility patterns are altered in NL3^R451C^ mice. **A, B, C, D:** Representative spatiotemporal heatmaps of caecal motor complexes (CaeMC) (denoted by black boxes and enlarged in **B, D**) in **A, B** wildtype (WT) and **C, D** NL3 mice. Forward contractions (FC) are denoted by black arrows and reverse contractions (RC) are denoted by dashed arrows. n=11 in each group. **E:** NL3 mice have higher number of CaeMC in 15 mins. **F:** Velocity of CaeMC between WT and NL3 is not significantly different but data shows an upward trend. **G:** Duration of CaeMC in NL3 are shorter compared to WT. **H:** No difference in quiescence period in WT and NL3 mice. n=10-11 in each group. n.s.=not significant. ns = p >0.05, *p <0.05.

### Caecal motility patterns are altered in NL3^R451C^ mice

Given that nNOS myenteric neuron populations were altered in the caecum (Sharna et al., 2020) and slower transit time was observed in NL3^R451C^ mice (Hosie et al., 2019), we assessed for potential changes in caecal motility in this model.

Using an inhouse software interface (as described in Swaminathan et al. (2016)), we characterised Caecal Motor Complexes (CaeMC) properties from video recordings. In the NL3^R451C^ mouse model of autism, the CaeMCs appear less complex than those observed in WT mice, with fewer forward and reverse patterns within each complex (**Fig 1B** wildtype, **1D** NL3). The frequency of CaeMCs occurring during 15 minutes of recording was higher in NL3^R451C^ mice compared to WT (**Fig 1E**, **Table 2**). In line with this observation, on average, each complex had a shorter duration in NL3^R451C^ compared to WT caecum (**Fig 1G**, **Table 2**). Analysis of CaeMC velocity showed a trend for an increased speed of contraction in the NL3^R451C^ caecum, however this was not statistically significant (**Fig 1F**, **Table 2**). Nevertheless, this trend for faster contractions is in line with the observed higher frequency and shorter duration of CaeMCs in mutant mice. The quiescence period (i.e., the period of rest between each CaeMC) was similar between WT and NL3^R451C^ (**Fig 1H, Table 2**).

**Table 2:** Analysis of CaeMCs including forward/pro-peristaltic and reverse/anti-peristaltic contraction parameters.

| Overall CaEMC parameters |  |  |  |  |  |
| --- | --- | --- | --- | --- | --- |
| Parameters | WT mean | NL3 mean | p-value | Benjamini-Hochberg Adjusted p-value | Significance |
| No. of CaEMC in 15min | 17.45 ± 1.0 | 24.09 ± 1.9 | 0.0103 | 0.0283 | * |
| CaEMC velocity (mm/s) | 1.48 ± 0.2 | 2.20 ± 0.3 | 0.037 | 0.0678 | NS |
| CaEMC duration (s) | 20.77 ± 2.4 | 11.59 ± 1.6 | 0.0048 | 0.0211 | * |
| CaEMC quiescence period (s) | 25.21 ± 2.1 | 26.21 ± 2.1 | 0.7363 | 0.8099 | NS |
| Percentage of F-R CaEMC (%) | 78.11 ± 2.7 | 55.73 ± 4.0 | 0.0001 | 0.0022 | ** |
| Caecal-colonic contraction interval (s) | 19.61 ± 2.2 | 8.476 ± 1.1 | 0.0002 | 0.0022 | ** |
| **Forward / Pro-peristaltic Contractions (FC)** |  |  |  |  |  |
| FC velocity (mm/s) | 4.12 ± 0.3 | 7.00 ± 1.1 | 0.0202 | 0.0494 | * |
| FC duration (s) | 3.80 ± 0.3 | 2.12 ± 0.2 | 0.0006 | 0.0044 | ** |
| FC Start position (%) | 92.17 ± 1.7 | 81.17 ± 2.6 | 0.0023 | 0.0127 | ** |
| **Reverse / Anti-peristaltic Contractions (RC)** |  |  |  |  |  |
| RC velocity (mm/s) | 7.94 ± 0.5 | 9.82 ± 1.3 | 0.1857 | 0.2724 | NS |
| RC duration (s) | 1.89 ± 0.1 | 1.37 ± 0.13 | 0.0082 | 0.0301 | * |

A canonical contraction pattern is observed within an individual CaeMC, where the forward contraction (FC) is followed by one or multiple reverse contractions (RC). The percentage of CaeMCs that possess this contraction pattern is significantly lower in NL3^R451C^ mice compared to WT (**Fig 2A, Table 2**). Some CaeMCs in both WT and NL3^R451C^ caecal preparations appear to lead to a subsequent contraction in the attached colon tissue. The caecal-colonic contraction interval, (i.e., the time elapsed between the start of the FC of the CaeMC and the start of a colonic contraction), was also altered with respect to genotype. In CaeMCs that precede a colonic contraction, the interval was shorter in NL3^R451C^ mice compared to WT (**Fig 2B, Table 2**). This finding could indicate that altered motility in the caecum is associated with downstream colonic dysmotility.

**Figure 2:**
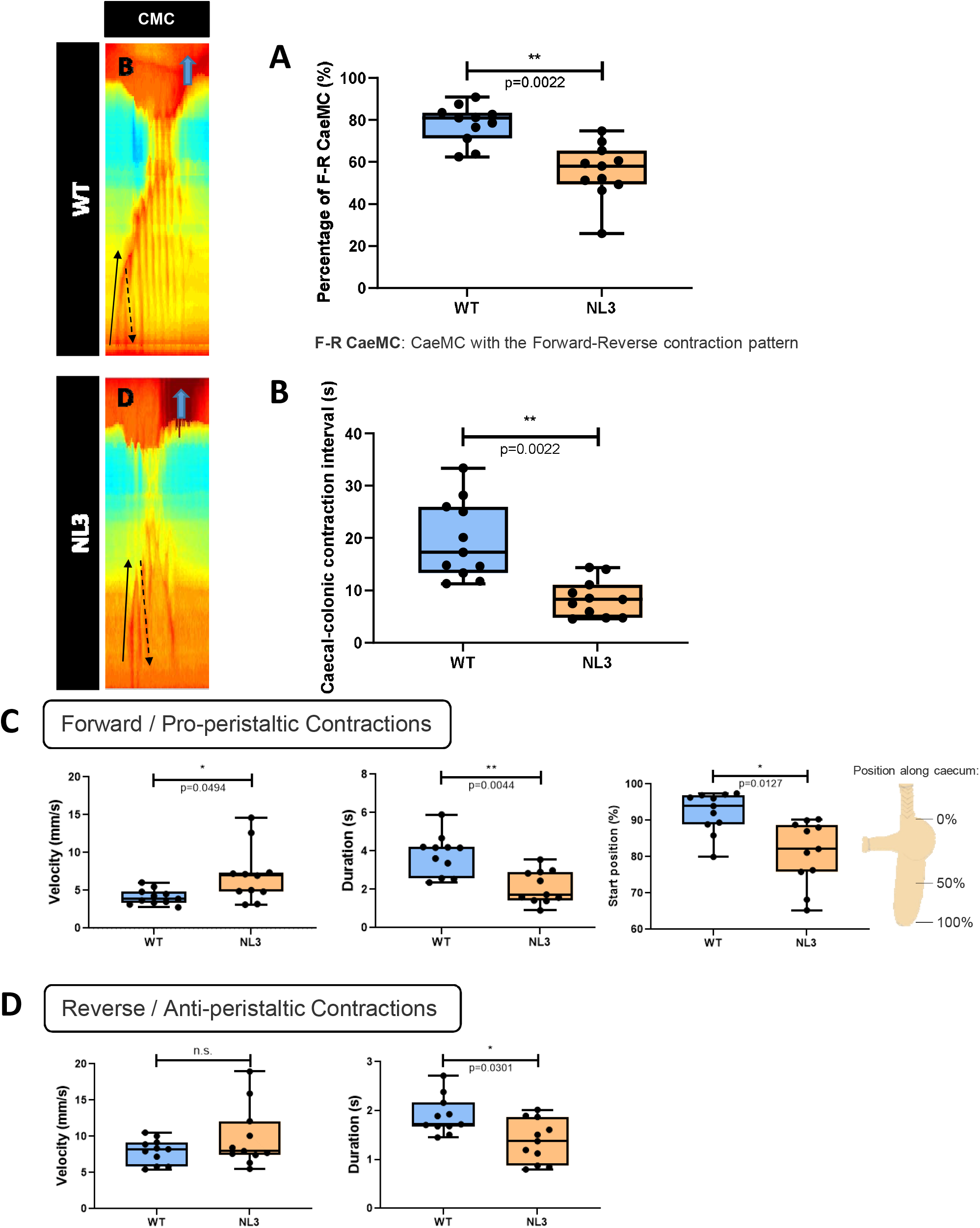
Altered caecal contraction patterns in NL3^R451C^ mice. **A:** The percentage of CaeMCs showing the contraction pattern is significantly lower in NL3 mice. **B:** the caecal-colonic contraction interval is significantly shorter in NL3 mice. n=11 in each group. ns = p >0.005, **p <0.01. Characteristics of individual contractions within CaeMCs. **C:** Velocity, duration, and start position of the forward contraction (FC) are altered in NL3 mice. **D:** In the reverse contraction (RC), the velocity is similar, but the duration is shorter in NL3 mice compared to WT. n=11 in each group. ns = p >0.05, *p <0.05, **p <0.01.

To better characterise caecal contraction patterns, CaeMCs were subsequently studied at a higher resolution to examine individual contractions within the contraction cluster. The first observed forward contraction (FC) of 3 CaeMCs for each video recording was chosen for analysis. These FCs had a higher velocity (**Fig 2C**) and shorter duration (**Fig 2C**) in NL3^R451C^ compared to WT mice (**Table 2**). The initiation position of the CaeMC was also altered in NL3^R451C^ mice. On average, the initiation of a CaeMC FC occurred mid-length of the caecum (i.e., closer to the colon) in NL3^R451C^ compared to WT caecum. In contrast, in WT samples, the FC started from the caecal tip and travelled a longer distance towards the colon (**Fig 2C**, **Table 2**). Reverse Contractions (RC) of CaeMCs were also studied. The first RC observed after the initial FC of 3 selected CaeMCs per mouse caecum was selected for detailed analysis. The RC in NL3^R451C^ mice occurred at a similar velocity to WT (**Fig 2D**) but had a significantly shorter mean duration (**Fig 2D**) in NL3^R451C^ compared to WT caecal samples (**Table 2**).

Overall, several properties of the caecal motility pattern in NL3^R451C^ mice were altered compared to WT littermates. CaeMCs occur more frequently and have a shorter duration in NL3^R451C^ mice. A lower percentage of CaeMCs showed the canonical FC-RC contraction pattern in NL3^R451C^ caecum. The interval between the start of a CaeMC and a contraction in the adjacent colon tissue was also reduced in NL3^R451C^ mice. FCs in NL3^R451C^ mice occurred at a higher velocity, persisted for a shorter duration, and were initiated at a mid-caecum location compared to a more distant caecal tip initiation in WT mice. RCs showed a similar trend to FCs in that a reduced contraction duration was observed in Nl3^R451C^ mice. These findings indicate the presence of caecal dysmotility in the NL3^R451C^ mouse model of autism.

### Caecal content is reduced but mucus content is unaffected in NL3^R451C^ mice

Reduced caecal weight was previously reported in NL3^R451C^ mice bred on a pure C57BL/6 background as well as mice generated on a mixed Sv129/C57BL/6 background (Sharna et al., 2020). A similar finding was also reported in NL3^−/-^ knockout mice, indicating that the *Nlgn3* gene plays a role in caecum physiology (Sharna et al., 2020). Although some caecal motility parameters are increased in NL3^R451C^ mice, these contractile patterns appear to be less efficient overall (more CaeMCs, shorter duration, fewer CaeMCs show a typical F-R contraction pattern and these travelled a shorter distance). Because the biofilm or mucus layer within the GI tract significantly contributes to gut permeability, the role of overall caecal content and mucus content in contributing to caecal weight difference was also investigated.

As previously reported, total caecal weight was lower in NL3^R451C^ mice compared to WT littermates (**Fig 3A**, **Table 3**). After clearing all content, caecal tissue was weighed again, and the weight of caecal content calculated. Similar to overall caecal weight, caecal content weight was also significantly lower in NL3^R451C^ mice compared to WT (**Fig 3B**, **Table 3**). In addition, when normalized against body weight, caecal content in NL3^R451C^ mice was significantly lower than WT littermates (**Fig 3C**, **Table 3**).

**Figure 3:**
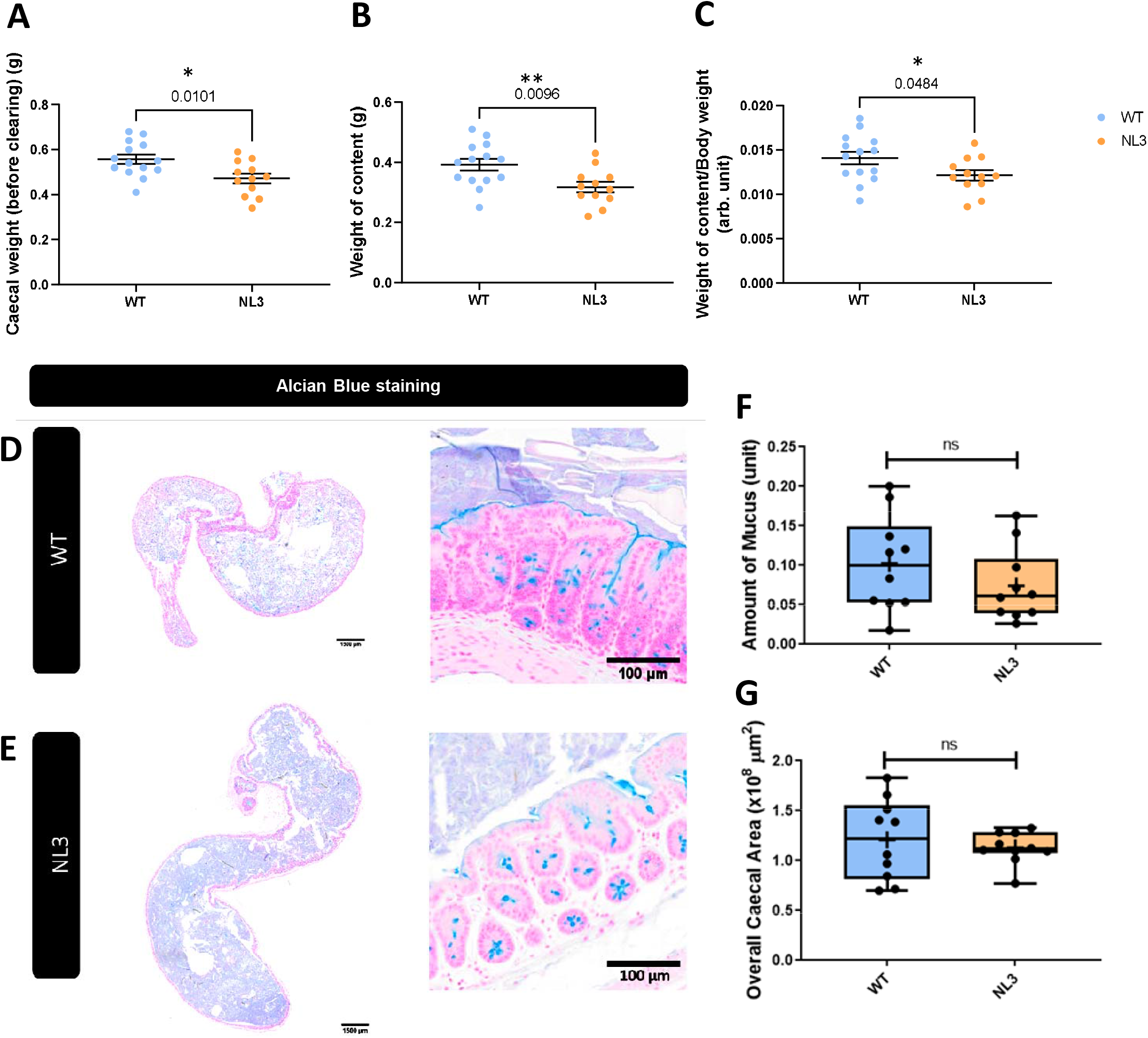
Caecal content is reduced in NL3^R451C^ mice. **A:** Caecal weight is decreased in NL3 mice. **B:** The weight of caecal content is reduced in NL3 mice compared to WT. **C:** The weight of caecal content after normalized to body weight is significantly lower in NL3 mice. WT, n=14; NL3, n=12. *p <0.05, **p <0.01. Amount of mucus content and overall caecal area in wildtype (WT) versus NL3 mice. Caecum cross-sections stained with Alcian Blue from WT (**D**) and NL3 (**E**) mice. **F**: There is no significant difference in amount of mucus in caecal cross-sections stained with Alcian Blue between WT and NL3 mice. **G**: There is no significant difference in overall caecal area between WT and NL3 mice. n=10 in each group. ns = p >0.05, *p <0.05, **p <0.01.

**Table 3:**
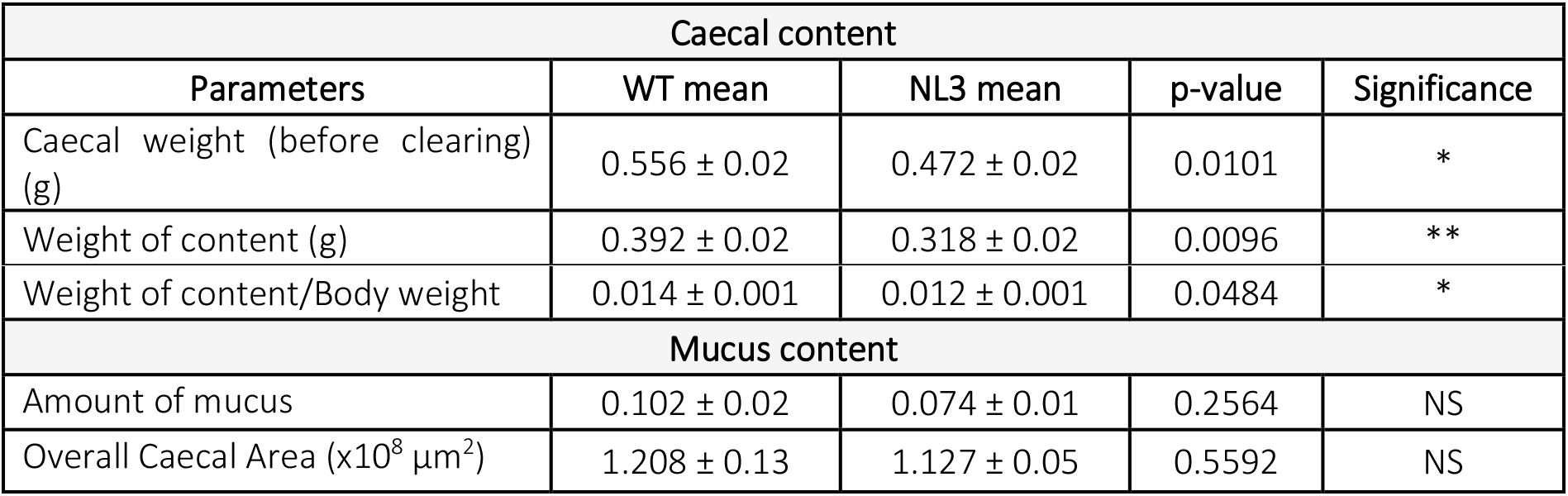
Analysis of caecal content and mucus content.

Mucus content was visualised in caecal cross sections using Alcian Blue staining and quantified using Image J software. There was no significant difference in the amount of mucus content observed in cross sections of caecum from both wildtype and NL3^R451C^ mice (**Fig 3D, 3E, 3F**, **Table 3**). The overall caecal area was also analysed as an index of caecum size to investigate if size correlated with caecal weight. Interestingly, caecal area was similar between wildtype and NL3^R451C^ mice (**Fig 3G**, **Table 3**).

### Unchanged gastrointestinal permeability alongside a subtle decrease in neurally-evoked secretion in the NL3^R451C^ caecum

The neuronal population in the submucosal plexus of the caecum in NL3^R451C^ mice was altered in that the number of nNOS neurons and total neurons were increased (Sharna et al., 2020). Since a primary function of the submucosal plexus layer of the ENS is to regulate gut secretion and permeability, and because ASD patients often present with increased permeability (commonly known as leaky gut syndrome), those functions were studied in WT and NL3^R451C^ caecum.

The transepithelial resistance (TER) was measured by monitoring the short-circuit current (I_SC_) in voltage-clamp mode. Caecal TER was similar in WT and NL3^R451C^ mice (**Fig 4A, Table 4**). The concentration of FITC that traversed the tissue preparation over time was also similar between WT and NL3^R451C^ caecal tissue samples (**Fig 4D**). In agreement with this finding, there was no difference in apparent permeability between the caecum from WT and mutant animals (**Fig 4B**) (**Table 4**). The TER over time was also measured throughout the duration of the permeability experiment to observe how TER changes over time and in response to FITC-Dextran. There was no significant difference between caecal TER in wildtype and NL3^R451C^ animals over time or in response to FITC-Dextran (**Fig 4C**, **Table 4**). There was, however, a slightly higher (but not statistically significant) increase in TER observed in NL3^R451C^ compared to WT animals (**Fig 4C**, **Table 4**).

**Figure 4:**
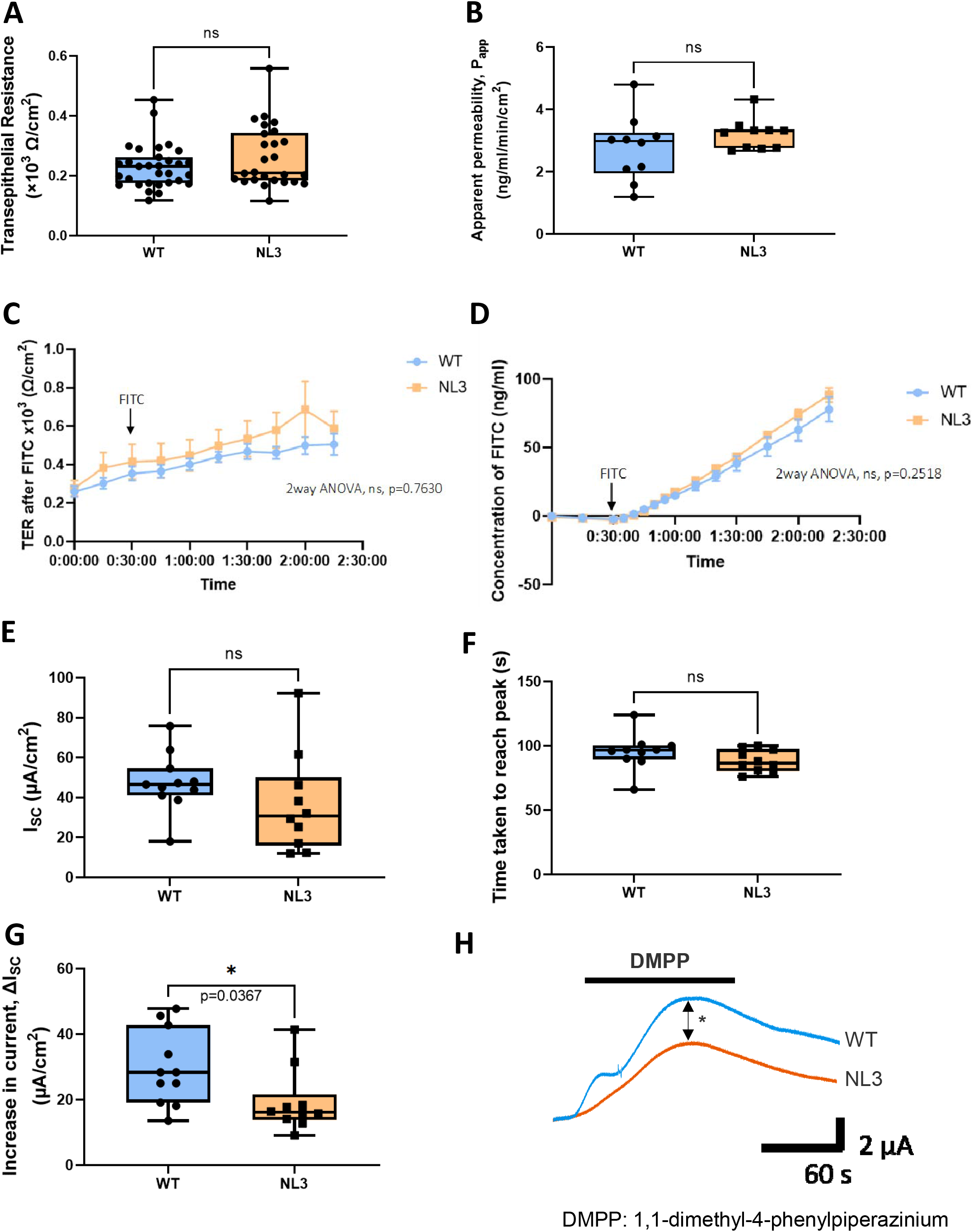
Transepithelial resistance (TER) and paracellular permeability of WT and NL3^R451C^ mice. There is no significant difference observed in the **A:** TER and **B:** apparent permeability of WT and NL3 caecum. The **D:** concentration of FITC-Dextran and **C:** TER over time did not show any difference in the caecum of WT and NL3 animals. A: WT: n=31; NL3: n=26. B, C, D: n=10 in each group. ns = p >0.005. Short-circuit current, I_SC_, response to 20 μM DMPP. **E**: The I_SC_ at baseline is not significantly different in NL3 compared to WT. **F**: Time taken to reach the peak of the trace is not altered in the caecum of NL3 mice compared to WT mice. **G**: The increase in current, **I** is significantly lower in the caecum of NL3 mice compared to

**Table 4:** Analysis of Ussing chamber parameters and response to 20 μM DMPP.

| Overall Ussing chamber parameters |  |  |  |  |
| --- | --- | --- | --- | --- |
| Parameters | WT mean | NL3 mean | p-value | Significance |
| Transepithelial resistance (TER) ($\times 10^3 \Omega/\text{cm}^2$) | 0.231 ± 0.01 | 0.262 ± 0.02 | 0.1809 | NS |
| TER after FITC ($\times 10^3 \Omega/\text{cm}^2$) | 0.406 ± 0.03 | 0.483 ± 0.04 | 0.7630 | NS |
| Concentration of FITC (ng/ml) | 21.14 ± 6.6 | 23.94 ± 7.7 | 0.2518 | NS |
| Apparent permeability, $P_{app}$ (ng/ml/min/$\text{cm}^2$) | 2.76 ± 0.3 | 3.20 ± 0.2 | 0.2400 | NS |
| Response to 20 μM DMPP | | | | |
| Short-circuit current, $I_{sc}$ ($\mu\text{A}/\text{cm}^2$) | 47.53 ± 4.4 | 36.66 ± 7.9 | 0.2320 | NS |
| Increase in current, $\Delta I_{sc}$ ($\mu\text{A}/\text{cm}^2$) | 29.78 ± 3.5 | 19.30 ± 3.1 | 0.0367 | * |
| Time taken to reach peak (s) | 95.4 ± 4 | 88.2 ± 3 | 0.1892 | NS |

Next, we investigated changes in short-circuit current (I_SC_) in response to DMPP application as an index of electrogenic secretion. Firstly, the I_SC_ at baseline showed no significant difference between WT and NL3^R451C^ caecum (**Supp. Fig 1A, Supp. Table 1**). Following the application of 10 μM DMPP (the nicotinic receptor antagonist), a similar increase in short-circuit current (ΔI_SC_) was observed in WT and NL3^R451C^ caecum (**Supp. Fig 1B, 1C**, **Supp. Table 1**). The time taken to reach the peak of the slope was also similar in both WT and NL3^R451C^ experimental groups (**Supp. Fig 1D, Supp. Table 2**).

Because the effects seen following application of 10 μM DMPP (i.e., an indication of neurally-evoked secretion) were subtle, another set of experiments was performed with a higher concentration of DMPP (i.e., 20 μM DMPP). Although the reduction of I_SC_ at baseline was not significantly different, there was a downward trend similar to that seen in the presence of 10 μM DMPP (**Fig 4E**, **Table 4**). In the presence of 20 μM DMPP, a significant decrease in the maximal I_SC_ response was observed in NL3^R451C^ compared to WT samples (**Fig 4G, 4H**, **Table 4**). The response time (i.e., the time taken to reach the peak of I_SC_ response) in the presence of 20 μM DMPP was also similar for both WT and NL3^R451C^ samples (**Fig 4F, Table 4**).

In conclusion, we report a subtle functional difference in fluid secretion in NL3^R451C^ caecum tissue where the peak I_SC_ was significantly lower in mutant mice compared to WT in response to DMPP application. Despite the clear differences in the number of neuronal subpopulations previously reported in the caecal submucosal plexus of NL3^R451C^ mice, only a small functional change in secretion was observed. When assessed, NL3^R451C^ and WT caecal preparations showed similar transepithelial resistance, TER and paracellular permeability across mucosa-submucosa preparations.

### VIP and ChAT submucosal neurons are not altered in the caecum of NL3^R451C^ mice

Vasoactive intestinal peptide (VIP) and acetylcholine (ACh) are key neurotransmitters of secretomotor neurons in the enteric nervous system involved in the neural control of secretion (Bornstein & Foong, 2018). Since subtle changes in neurally-evoked secretion were observed in the caecum of NL3^R451C^ mice, submucosal neurons were stained for these major neurotransmitters to increase understanding of biological mechanisms behind the observed changes in the response to DMPP. First, the total number of neurons per ganglion were counted to gauge ganglia size. In contrast with our previously published findings (Sharna et al., 2020), we did not observe an increase in the total number of neurons per ganglion in the submucosal plexus of NL3^R451C^ caecum compared to WT mice in the current study (**Fig 5C**, **Table 5**).

**Figure 5:**
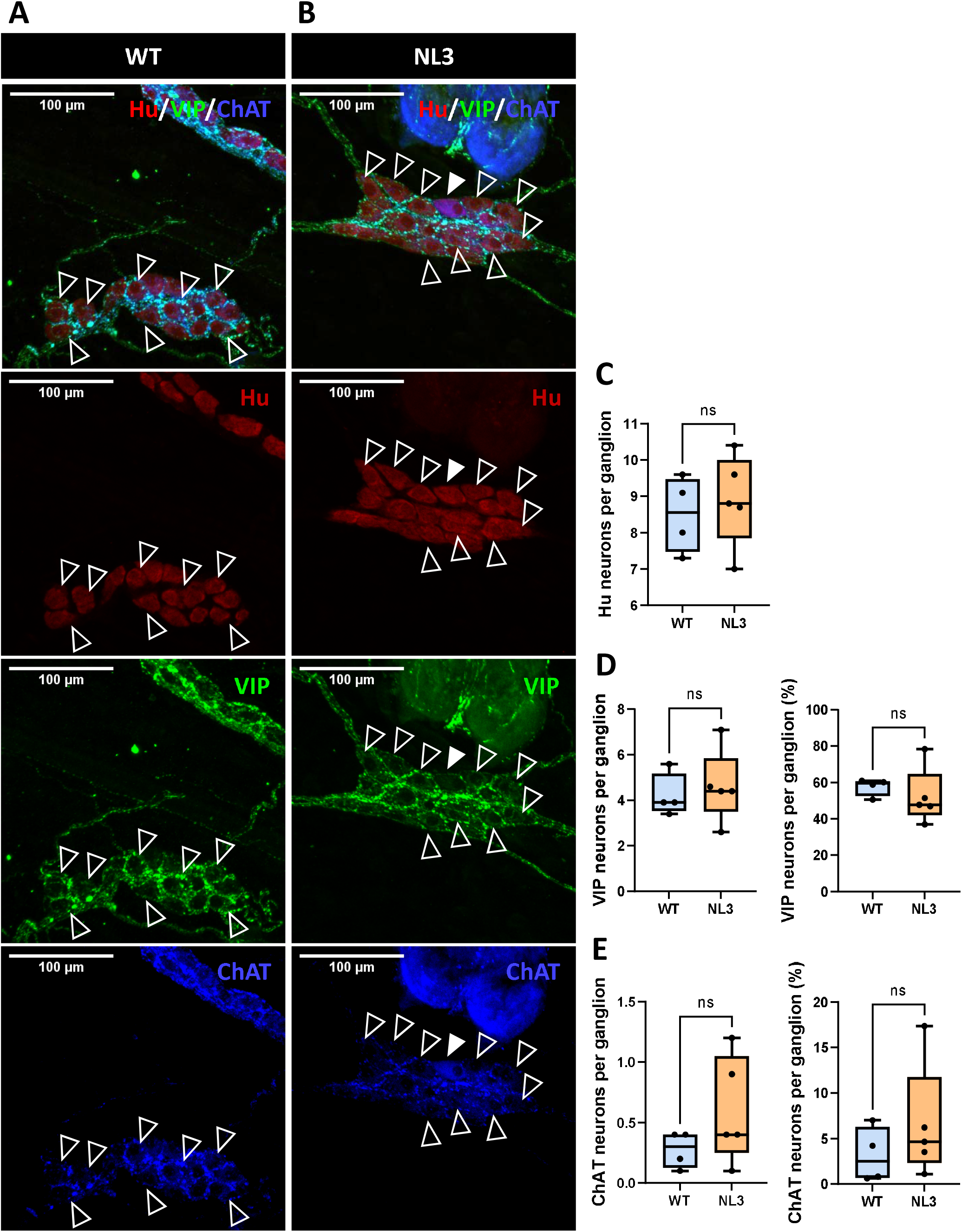
Hu, VIP, and ChAT staining in the submucosal plexus of WT (**A**) and NL3^R451C^ (**B**) mice. VIP (clear arrowheads) and ChAT (filled arrowhead) enteric neurons were visualized in the caecal submucosal plexus using immunostaining. **C**: No significant difference was observed in the number of Hu neurons per ganglion. **D**: The number of VIP stained neurons and the percentage of VIP neurons per ganglion also showed no significant difference between WT and NL3 mice. **E**: No significant difference was observed in the number of ChAT neurons and the percentage of ChAT neurons per ganglion. n=4-5 in each group. WT: n=4; NL3: n=5; ns = p >0.05.

**Table 5:** Analysis of Hu, VIP, and ChAT-stained submucosal neurons.

| Submucosal Hu<sup>+</sup> neurons |  |  |  |  |
| --- | --- | --- | --- | --- |
| Parameters | WT mean | NL3 mean | p-value | Significance |
| Hu neurons per ganglion | 8.50 ± 0.5 | 8.90 ± 0.6 | 0.6273 | NS |
| Submucosal VIP<sup>+</sup> neurons |  |  |  |  |
| VIP neurons per ganglion | 4.20 ± 0.5 | 4.62 ± 0.7 | 0.6611 | NS |
| VIP neurons per ganglion (%) | 57.64 ± 2.4 | 52.31 ± 7.0 | 0.5337 | NS |
| Submucosal ChAT<sup>+</sup> neurons |  |  |  |  |
| ChAT neurons per ganglion | 0.28 ± 0.1 | 0.60 ± 0.2 | 0.2064 | NS |
| ChAT neurons per ganglion (%) | 3.16 ± 1.5 | 6.57 ± 2.8 | 0.3589 | NS |

**Table 6:**
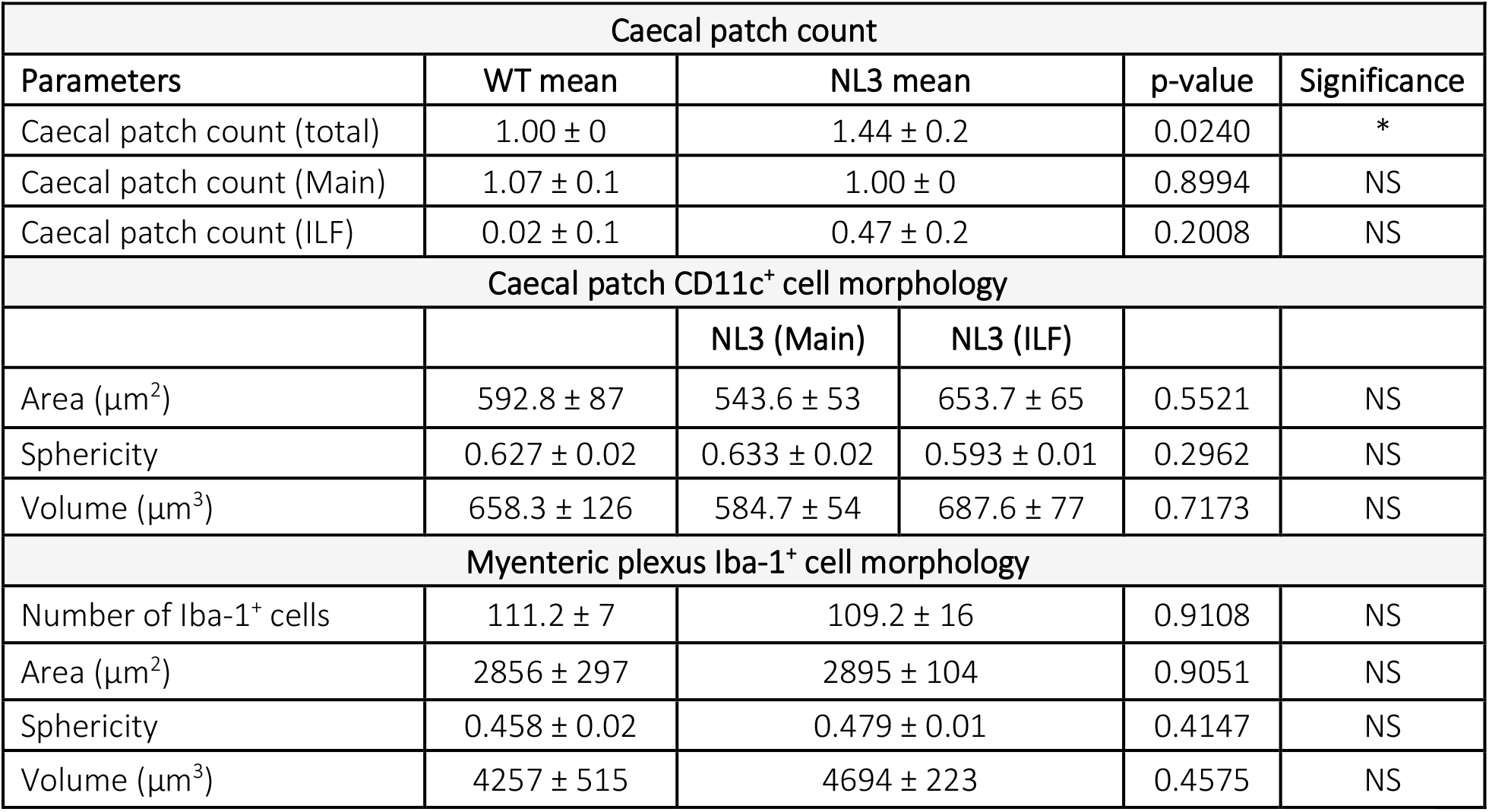
Analysis of caecal patch count, CD11c^+^ cell morphology and Iba-1^+^ cell morphology.

| Caecal patch count |  |  |  |  |  |
| --- | --- | --- | --- | --- | --- |
| Parameters | WT mean | NL3 mean |  | p-value | Significance |
| Caecal patch count (total) | 1.00 ± 0 | 1.44 ± 0.2 |  | 0.0240 | * |
| Caecal patch count (Main) | 1.07 ± 0.1 | 1.00 ± 0 |  | 0.8994 | NS |
| Caecal patch count (ILF) | 0.02 ± 0.1 | 0.47 ± 0.2 |  | 0.2008 | NS |
| Caecal patch CD11c<sup>+</sup> cell morphology |  |  |  |  |  |
|  |  | NL3 (Main) | NL3 (ILF) |  |  |
| Area (µm<sup>2</sup>) | 592.8 ± 87 | 543.6 ± 53 | 653.7 ± 65 | 0.5521 | NS |
| Sphericity | 0.627 ± 0.02 | 0.633 ± 0.02 | 0.593 ± 0.01 | 0.2962 | NS |
| Volume (µm<sup>3</sup>) | 658.3 ± 126 | 584.7 ± 54 | 687.6 ± 77 | 0.7173 | NS |
| Myenteric plexus Iba-1<sup>+</sup> cell morphology |  |  |  |  |  |
| Number of Iba-1<sup>+</sup> cells | 111.2 ± 7 | 109.2 ± 16 |  | 0.9108 | NS |
| Area (µm<sup>2</sup>) | 2856 ± 297 | 2895 ± 104 |  | 0.9051 | NS |
| Sphericity | 0.458 ± 0.02 | 0.479 ± 0.01 |  | 0.4147 | NS |
| Volume (µm<sup>3</sup>) | 4257 ± 515 | 4694 ± 223 |  | 0.4575 | NS |

We also stained for VIP and ChAT in the submucosal plexus to identify if subpopulations of the two key neurochemically distinct neuronal populations have been altered in mutant mice. The ChAT enzyme is directly involved in acetylcholine synthesis and therefore ChAT^+^ neurons encode for cholinergic neurons. Another key population of neurons are non-cholinergic neurons which express VIP but do not possess ChAT. In the submucosal plexus of WT and NL3^R451C^ caecum, the numbers and proportions of VIP (**Fig 5D**) and ChAT (**Fig 5E**) neurons per ganglion were similar (**Table 5**).

### Altered gut-associated lymphoid tissue (caecal patches) in NL3^R451C^ caecum

A regional specific aggregate of gut-associated lymphoid tissue called the caecal patch is typically found at the blind end of the caecum. Previously, Sharna et al. (2020) reported that gut macrophages within this lymphoid tissue demonstrated altered morphology, suggesting a reactive immune phenotype. Here we studied the gross morphology of the caecal patch (**Fig 6A, 6B**) and revealed a higher number of total caecal patches in NL3^R451C^ caecum compared to WT littermates (**Fig 6C**, **Table 5**). There is typically a single primary caecal patch located at the tip of the caecal pouch, however additional patches or isolated lymphoid follicles (ILF) are more commonly observed in the NL3^R451C^ caecum. These additional patches are referred to as ILF due to their smaller size that likely indicates they contain a single germinal centre with an overlying layer of follicle-associated epithelium. These additional caecal patches or ILFs are smaller in size and are generally located anterior to the mesenteric border in-line with the main patch along the body of the caecum (**Fig 6B**).

**Figure 6:**
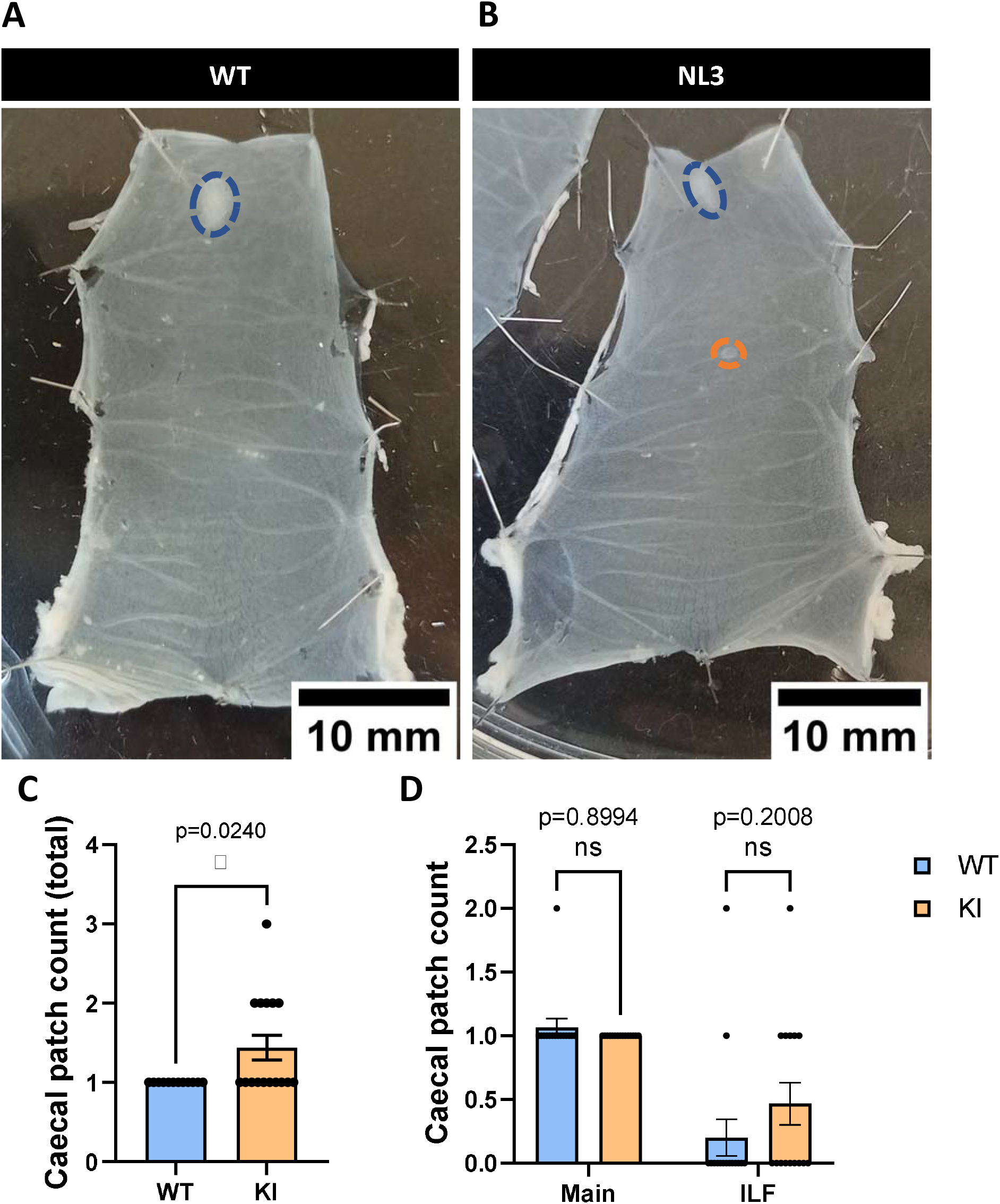
Multiple caecal patches are present in NL3^R451C^ mice. Representative images of pinned-out caecum from WT (**A**) and NL3 (**B**) mice. Blue dashes outline the main caecal patch, whereas orange dashes outline the isolated lymphoid follicles (ILF) typically found along the body of the caecum. **C:** Total caecal patch is significantly higher in NL3 caecum than WT littermates. 3 outliers were removed in the WT group according to the ROUT Q=1% method. **D:** When total caecal patch count is split into main and ILF caecal patches, the difference is not significant albeit an increasing trend in NL3 ILF. WT: n=12; NL3: n=16. ns = p >0.05, *p <0.05.

To investigate for morphological changes in immune cells as an indication of altered immune activity, caecal patch cross sections were stained for the dendritic cell marker, CD11c. CD11c is present on many immune cells including macrophages, but it is a known marker for dendritic cells. Antigen-presenting dendritic cells localized in the GALT along the gastrointestinal tract play an important role in IgA synthesis in response to commensal bacteria within the lumen (Tezuka & Ohteki, 2019). Given that the number of caecal patches observed in NL3^R451C^ mice are increased, we investigated whether the CD11c^+^ cellular population located within caecal patches is also altered.

Only main caecal patch tissues were processed for the WT group (**Fig 7A**), whereas both main and ILF caecal patches were processed for the NL3^R451C^ group (**Fig 7B, 7C**). 3-dimensional reconstruction and analysis of CD11c^+^ cells of WT and NL3^R451C^ caecal patch tissue, however, showed no morphological differences in cellular area (**Fig 7D**), sphericity (**Fig 7E**), or volume (**Fig 7F**) (**Table 5**).

**Figure 7:**
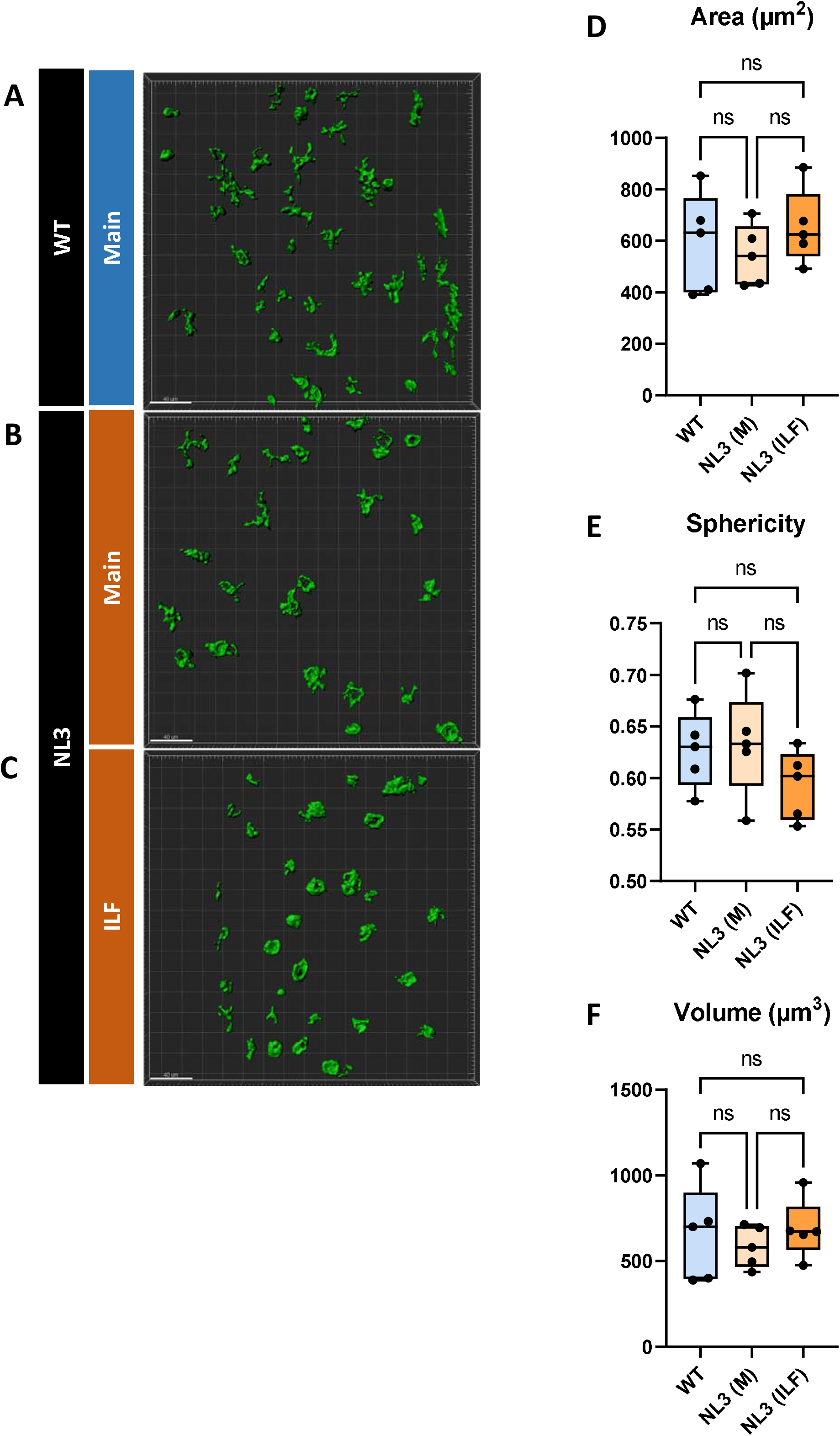
No morphological difference observed in CD11c^+^ cells in caecal patch of NL3^R451C^ and WT mice. Both main and ILF NL3 caecal patches (**A, B**) were stained for CD11c whereas only main patches were stained for WT mice (**C**). No significant morphological differences were observed among WT main, NL3 main, and NL3 ILF caecal patches with the parameters area (**D**), sphericity (**E**), and volume (**F**). n=5 in each group. ns = p >0.005.

Since Iba-1^+^ gut macrophages within NL3^R451C^ caecal patch samples showed morphological differences suggestive of a reactive phenotype (Sharna et al., 2020), Iba-1^+^ macrophages were investigated within the myenteric plexus layer of WT and NL3^R451C^ caecal tissue (**Fig 8A, 8B**). There was no difference in the number of Iba-1^+^ cells in the myenteric plexus of NL3^R451C^ caecum compared to WT littermates (**Fig 8C**, **Table 5**). 3D analysis of Iba-1^+^ gut macrophages also showed no morphological difference in area (**Fig 8D**), sphericity (**Fig 8E**) or volume (**Fig 8F**) in the caecal myenteric plexus of NL3^R451C^ compared to WT littermate mice (**Table 5**).

**Figure 8:**
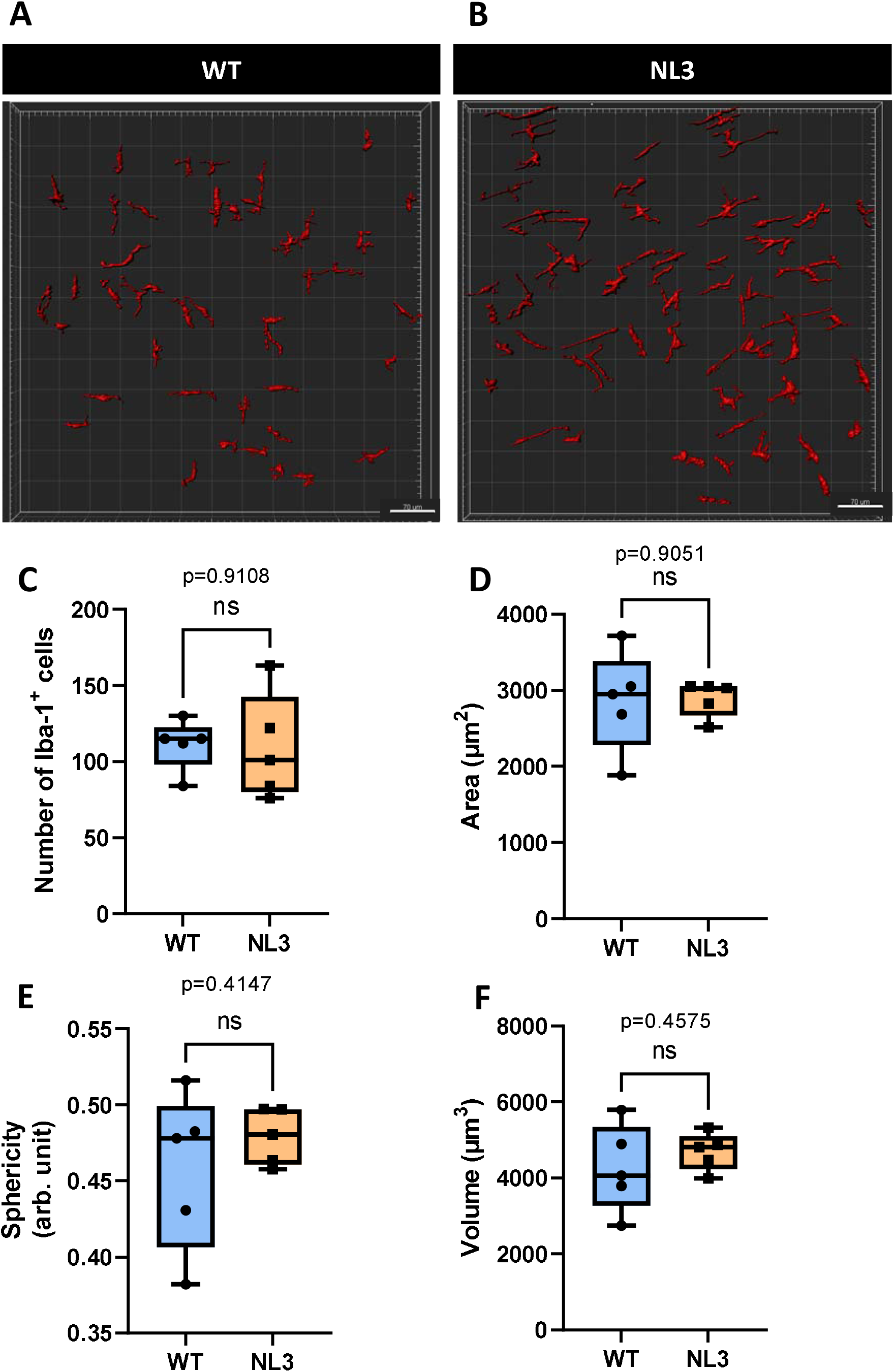
No morphological difference was observed in Iba-1^+^ cells in the myenteric plexus of NL3^R451C^ caecum compared to WT littermates. 3D reconstruction and analysis of Iba-1^+^ cells in WT (**A**) and NL3 (**B**) myenteric plexus. No significant morphological differences were observed with the parameters number of Iba-1^+^ cells (**C**), area (**D**), sphericity (**E**), and volume (**F**). n=5 in each group. ns = p >0.05.

## Discussion

Gastrointestinal symptoms are often reported in patients with neurodevelopmental disorders (Gorrindo et al., 2012; McElhanon et al., 2014; Restrepo et al., 2020; Thulasi et al., 2019), with dysmotility likely being an underlying factor. This has been mirrored in animal models of neurodevelopmental disorders, whereby visualised motility patterns are often altered (Fröhlich et al., 2019; Hosie et al., 2019). Alterations in caecal weight and microbial populations are common findings in a myriad of animal models including germ-free (Smith et al., 2007), and antibiotic-treated (Reikvam et al., 2011) rodents and in models of inflammation and neurological diseases (Houlden et al., 2016; Sharna et al., 2020), but these observations are often not pursued further.

Here, we report that caecum motility is altered in the NL3^R451C^ mouse model of autism, including an increase in CaeMC frequency alongside shorter CaeMC durations. We speculate that this dysmotility phenotype decreases the efficiency of contractile activity in NL3^R451C^ mice and contributes to a reduced volume of luminal content within the caecum, leading to a decrease in caecal weight as previously reported (Sharna et al., 2020). To better understand the role of the caecum, it is crucial to characterise caecal motility as a measure of caecal dysfunction in preclinical models as well as potential alterations in caecal weight. In the current study, we observed that mouse CaeMCs predominantly consisted of multidirectional contractions, whereas multidirectional contraction clusters did not appear to be the dominant pattern in the rabbit caecum (Hulls et al., 2016). Should caecal dysfunction be identified across animal models (for example, preclinical models of neurodevelopmental disorders) this location could provide a target for therapeutic design to enhance gut health.

Based on the observed changes in Caecal Motor Complexes (CaeMCs) in the NL3^R451C^ caecum, caecal motility in NL3^R451C^ mice appears to be less effective compared to WT mice. This could lead to alterations in the churning and mixing motion of the caecum resulting in impaired fermentative digestion. While enzymatic digestion and absorption of nutrients is the main function of most regions along the gastrointestinal tract, the caecum is the primary site for fermentation (Girard-Madoux et al., 2018; Laurin et al., 2011; Oltmer & Engelhardt, 1994). Fermentation of chyme in the caecum is a slower process compared to enzymatic digestion and the end-product is rich in nourishing metabolites such as short chain fatty acids (SCFA) (Brown et al., 2018). This slower fermentation process requires the chyme to stay in the caecum for a longer period; hence a slower motility profile may be required compared to other regions of the gastrointestinal tract (Hulls et al., 2012). The faster motility observed in the NL3^R451C^ caecum could result in incomplete fermentation of the luminal content. The lack of the canonical Forward-Reverse contraction pattern within CaeMCs in the NL3^R451C^ caecum could also indicate that the churning of luminal contents is less effective. These observations could negatively impact the fermentation process and thus the production of SCFA. Fermentation products such as SCFA from carbohydrates are crucial in maintaining a nourishing and enriched environment for host colonocytes that are positioned directly distal to the caecum. From a clinical perspective, specific gut microbiota-derived metabolites are strongly correlated with ASD clinical behavioural scores (Needham, Adame, et al., 2020). Strategies to alter metabolite levels by “replenishing” or administering SCFAs typically found at low levels in patient profiles have been utilized to induce a rescue effect in models of ASD, Alzheimer’s disease, and Huntington disease (Gardian et al., 2005; Kidd & Schneider, 2010; Kilgore et al., 2010; Needham, Kaddurah-Daouk, et al., 2020; Sharon et al., 2019; Vuong et al., 2020). Based on these findings and the changes observed in the current study, fermentation of digesta in the caecum could be key to supplementing the microbiome with the SCFAs that are critical for a healthy gut environment.

In this study, we highlighted that caecal dysmotility in NL3^R451C^ mice impacted downstream contractions in the proximal colon. Due to size limitations of the organ bath, the ileo-caecal-colonic preparation dissected for caecal motility experiments included the proximal colon region only. It is therefore not known whether these proximal colonic contractions have the potential to migrate to more than half the length of the colon and could therefore be considered a colonic migrating motor complex (CMMC) (Fida et al., 1997). Nevertheless, the caecal-colonic contraction interval (i.e., the time elapsed between the start of the CaeMC and the start of a colonic contraction) is shorter in NL3^R451C^ mice. This suggests that colonic function, such as water absorption and fecal pellet formation, could be affected by a motility defect in the caecum, an upstream (proximal) segment of the gastrointestinal tract.

In contrast with clinical reports identifying an association of “leaky gut” conditions with ASD, no significant difference was reported in the transepithelial resistance or permeability of the caecum in WT and NL3^R451C^ mice. Subtle changes permeability could be revealed however if NL3^R451C^ mice were studied in a fasted state, since gut permeability is known to be modified in response to food intake and circadian rhythms (Seillet et al., 2020; Talbot et al., 2020). Although caecal permeability is unchanged in non-fasted NL3^R451C^ mice, altered permeability may occur in other segments of the gastrointestinal tract in this model given the vast differences in specialized function of different GI regions. Such potential changes could be determined by studying the gastrointestinal tract permeability of NL3^R451C^ mice in a region-specific manner. Basal short-circuit current remained unchanged in NL3^R451C^ mice, indicating that at resting physiological conditions, mucosal secretion is not affected by the NL3^R451C^ mutation at neuronal synapses. When submucosal neurons were selectively stimulated with DMPP, however, NL3^R451C^ caecal tissue showed a significantly reduced peak secretory response compared to WT. This finding suggests that the NL3^R451C^ mutation alters nicotinic acetylcholine receptor (nAChR) expression given that DMPP acts exclusively on this receptor subtype. The intestinal mucosal secretory system plays a vital function in secreting water and mucus to flush foreign pathogens out of the gastrointestinal lumen. Decreased secretory function in the NL3^R451C^ caecum could indicate a defective pathogen removal system. When the main subpopulations of the submucosal plexus were investigated, however, we found no significant difference in the proportions of ChAT and VIP-stained secretomotor neurons. Nevertheless, there was a trend for increased numbers of ChAT neurons in NL3^R451C^ mice which may explain the subtle abnormality in the response to DMPP observed in Ussing chamber experiments in NL3^R451C^ caecal tissue. These findings suggest a plausible mechanism by which an autism-associated mutation that impacts synaptic function might alter gastrointestinal secretory function via a nAChR-mediated pathway in the submucosal plexus.

The human appendix is often referred to as an immune-rich organ because of the significant amount of lymphoid tissue present at this location. Similarly, the mouse caecum typically possesses one prominent lymphoid tissue aggregate at the apex of the blind-ended pouch. In this study, significantly more caecal patches were found in the caeca of NL3^R451C^ mice compared to WT littermates. In instances where more than one caecal patch was observed, the patches were noticeably smaller, positioned anterior to the mesenteric border along the body of the caecum, and are thus referred to as isolated lymphoid follicles (ILF). Masahata et al. (2014) previously described the function of IgA-secreting cells from both caecal and Peyer’s patches, highlighting that while there is some overlap in function, IgA^+^ cells that migrate to the colon from the caecal patch can alter microbiota composition whereas this was not described for similar cells located in Peyer’s patches. Based on our findings of increased caecal patches, it is proposed that both caecal microbiota composition and function are altered.

We aimed to investigate the immune activity of caecal patch tissue to understand how an increase in number of patches observed might affect immune function. CD11c was selected as a dendritic cell marker to investigate the microbe-sensing ability of the patches (Everett et al., 2004). There was no morphological difference observed in the area, volume, and sphericity of the cells between caecal patches from WT and NL3^R451C^ mice. Since previous findings from Sharna et al. (2020) found that NL3^R451C^ gut macrophages in the caecal patch were rounder, present in higher density, and had a smaller volume, Iba-1 was used as a pan-macrophage marker to investigate if a similar profile is found in myenteric gut macrophages. Similar to CD11c^+^ cells, no morphological differences were observed in Iba-1^+^ stained cells in the myenteric plexus of NL3^R451C^ caecum. This negative finding highlights potential specificity in neuroimmune modification whereby only certain populations of immune cells (i.e., Iba-1 expressing macrophages) are altered in the caecal patch compared to other immune cells (CD11c^+^) or even in other regions of the caecum (i.e. within the enteric nervous system layers).

The NL3^R451C^ genetic mouse model of autism is a well-studied model of central nervous system dysfunction. Recent additions to the literature have shown that changes to the NL3 post-synaptic protein also influence enteric nervous system activity (Hosie et al., 2019; Leembruggen et al., 2019; Sharna et al., 2020). Here, we show that ASD-associated gastrointestinal dysfunction impacts not only the colon and small intestine, but the caecum as well. The caecum is an important focal point for studies on neuro-immune-microbe interactions due to the presence of an intrinsic network of neurons within the caecal walls, a rich profile of immune cells and lymphoid tissue, and its proposed function as reservoir for commensal microbiota. To date, the Neuroligin-3 protein has not been identified in gastrointestinal lymphoid tissue or specific enteric neurons. This study provides evidence that the NL3^R451C^ mutation impacts gastrointestinal function, specifically caecal dysmotility, secretion, and gut-associated lymphoid tissue structure. These observations link an ASD-associated missense mutation in the gene encoding Neuroligin-3 to caecal dysfunction, which could influence luminal microbial composition, and, as a consequence, digestive fermentation and microbial metabolite production in individuals expressing this gene mutation. Whether this phenomenon applies to other ASD-associated mutations affecting nervous system function (for example, Shank3 deletion), remains unknown. If multiple ASD-relevant mutations impacting the nervous system also cause gastrointestinal dysfunction such as the caecal changes we report here, this may point to a broader biological mechanism contributing to gut symptoms in individuals with ASD. Furthermore, microbial metabolites are a key mediator in the cross-talk between the gut and brain and are therefore likely important for maintaining interactions between microbes and the enteric nervous system. Certain microbial metabolites, for example, are liposoluble and can cross the blood brain barrier to interact with neurons and immune cells of the central nervous system. In conclusion, targeting dysfunction in the caecum could assist with potential therapeutic strategies to alter metabolite profiles and improve gastrointestinal symptoms in ASD.

## Acknowledgements

This project was support by the NHMRC Ideas Grant APP2003848 “Identifying how the enteric nervous system regulates gut permeability in autism” awarded to E.L.H-Y. C.Y.Q.L. received the RMIT Research Stipend Scholarship (RRSS). No role of funder in design, analysis or reporting of the study.

## Author contributions

C.Y.Q.L. performed all experiments, interpreted the data, and drafted the manuscript. G.K.B. and M.H. were involved in and assisted with the *ex vivo* caecal motility assay and Ussing chamber experiments respectively. A.E.F. and E.L.H-Y. conceptualize and designed the study. All authors approved the manuscript for submission.

---

**Supplementary Video 1:**
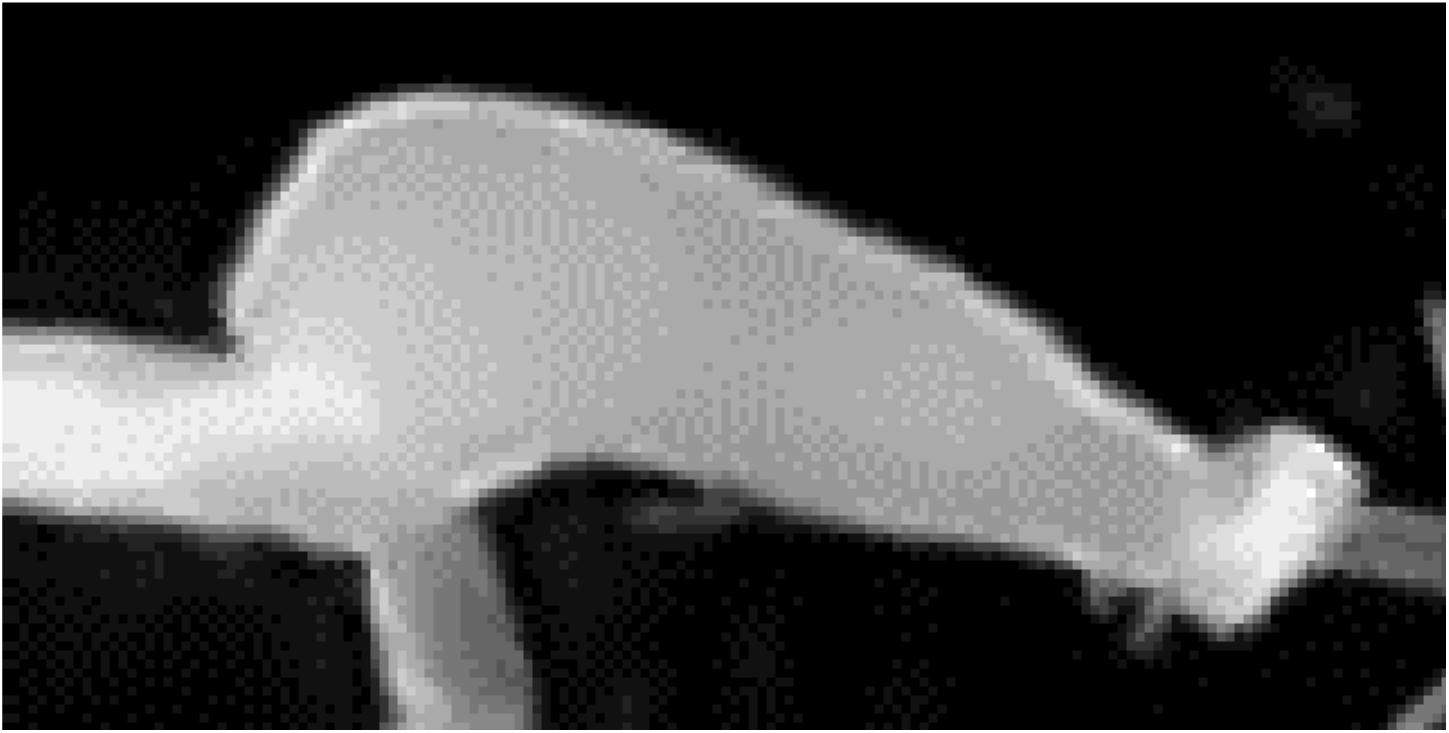
A real time video recording of contractions in the caecum.

**Supplementary Figure 1:**
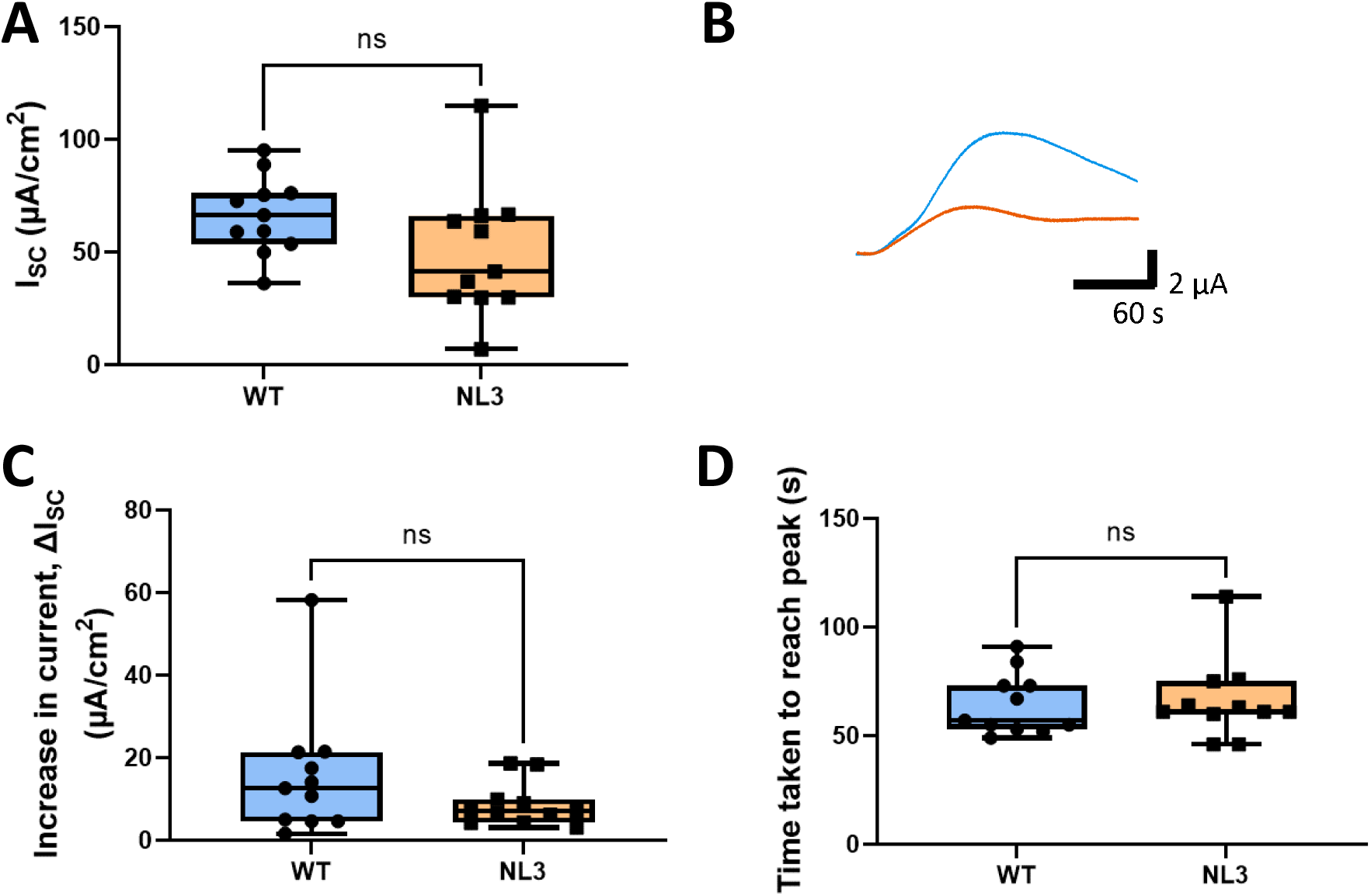
Short-circuit current, I_SC_, response to 10 μM DMPP. **A**: The I_SC_ at baseline is not significantly different in NL3^R451C^ compared to WT. **B**: Representative current traces versus time of increase in I_SC_ in WT (blue) and NL3 (orange). **C**: The increase in current, ΔI_SC_ and **D**: time taken to reach the peak of the trace is not altered in the caecum of NL3 mice compared to WT mice. n=11 in each group. ns = p >0.05.

**Supplementary Table 1:** Analysis of caecal tissue response to 10 μM DMPP.

| Response to 10 $\mu M$ DMPP | | | | |
| --- | --- | --- | --- | --- |
| Short-circuit current, $I_{sc}$ ($\mu A/cm^2$) | $66.56 \pm 5.2$ | $49.57 \pm 8.7$ | 0.1097 | NS |
| Increase in current, $\Delta I_{sc}$ ($\mu A/cm^2$) | $15.64 \pm 4.7$ | $8.636 \pm 1.6$ | 0.1770 | NS |
| Time taken to reach peak (s) | $64.4 \pm 4$ | $66.1 \pm 6$ | 0.8178 | NS |

